# Mathematical modelling of autoimmune myocarditis and the effects of immune checkpoint inhibitors

**DOI:** 10.1101/2021.09.03.458857

**Authors:** Solveig A. van der Vegt, Liudmila Polonchuk, Ken Wang, Sarah L. Waters, Ruth E. Baker

**Affiliations:** Wolfson Centre for Mathematical Biology, Mathematical Institute, University of Oxford, Oxford, UK; Roche Pharmaceutical Research and Early Development, Roche Innovation Center Basel, F. Hoffmann-La Roche Ltd., Basel, Switzerland; Oxford Centre for Industrial and Applied Mathematics, Mathematical Institute, University of Oxford, UK

**Keywords:** myocarditis, autoimmunity, mathematical modelling, immune checkpoint inhibitors

## Abstract

Autoimmune myocarditis is a rare, but frequently fatal, side effect of immune checkpoint inhibitors (ICIs), a class of cancer therapies. Despite extensive experimental work on the causes, development and progression of this disease, much still remains unknown about the importance of the different immunological pathways involved. We present a mathematical model of autoimmune myocarditis and the effects of ICIs on its development and progression to either resolution or chronic inflammation. From this, we gain a better understanding of the role of immune cells, cytokines and other components of the immune system in driving the cardiotoxicity of ICIs. We parameterise the model using existing data from the literature, and show that qualitative model behaviour is consistent with disease characteristics seen in patients in an ICI-free context. The bifurcation structures of the model show how the presence of ICIs increases the risk of developing autoimmune myocarditis. This predictive modelling approach is a first step towards determining treatment regimens that balance the benefits of treating cancer with the risk of developing autoimmune myocarditis.

## 1. Introduction

Immune-related adverse events, and autoimmunity in particular, are an extremely common side-effect of immune checkpoint inhibitors (ICIs), a class of treatments for which approximately 40% of US cancer patients are eligible [1]. Autoimmune side-effects are experienced by 86-96% of cancer patients treated with ICIs, and in 17-59% of patients these side-effects are severe to life-threatening [2, 3]. Myocarditis, inflammation of cardiac muscle tissue, is a rare but frequently fatal side effect of ICIs. Of patients treated with ICIs, about 0.1–1% will develop myocarditis, which is a significant increase from the 0.022% incidence rate in the general population [4, 5]. Furthermore, myocarditis proves fatal in 25–50% of cases [6, 7]. In common with other inflammatory diseases such as rheumatoid arthritis and asthma, the pathology comes from the failure of inflammation to resolve itself, instead becoming a chronic condition [8]. In autoimmune myocarditis, this chronic stage is driven by CD4+ T cells and leads to irreversible tissue damage and possibly heart failure [9, 10].

Despite significant experimental research devoted to gaining an improved understanding of autoimmune myocarditis, the immunological mechanisms underlying the development and progression of this disease are not well understood [6, 9]. The immense complexity of the immune system makes it difficult the determine which cell types and pathways are essential, and which are of secondary importance. Making this distinction is necessary for moving research in this area forward in an efficient manner. Furthermore, no pre-clinical *in vitro* cardiotoxicity assay to asses whether potential new ICIs might increase the risk of autoimmune myocarditis currently exists. Development of such an assay will depend on the identification of the cell types that play a critical role in the disease.

Although mathematical modelling has been extensively applied to better understand the progression of diseases including cancer [11, 12, 13], neurological disorders [14, 15, 16], and autoimmune conditions [17, 18], to name but a few, no mathematical model of autoimmune myocarditis exists to our knowledge. Mathematical modelling provides a framework to deal with the highly complex, nonlinear interactions of the immune system, and can increase understanding of the mechanisms underlying ICI-induced autoimmune myocarditis. As such, the aim of this work is to develop the first mathematical model of autoimmune myocarditis, which will aid in determining which immunological pathways are essential to its development and progression, and which are of secondary importance.

Existing mathematical models of the immune response provide a starting point for the modelling of autoimmune myocarditis, but none suffice on their own to describe the specific mechanisms of this disease. Many, such as those consisting of partial differential equation or agent-based models, are currently too complex for purpose. Ordinary differential equation (ODE)-based models, such as Sontag’s model of a generalized immune response [19], lack spatial effects and can be used to describe the autoimmune cycle in an analytically tractable way. Carneiro et al. [20] present a ODE model that zooms in on the clonal expansion of T cells and self-tolerance, two important processes in the development of autoimmunity in autoimmune myocarditis. More detail quickly leads to more complex mathematics, as seen in the model of T-cell mediated autoimmunity by Delitala et al. [21], which consists of both ODEs and integro-differential equations. Su et al. [22] develop a PDE model that describes the minimally required set of immune cells to simulate the immune system. These more detailed models [21, 20, 22] lack the mathematical simplicity that is desirable in a first model of autoimmune myocarditis, and none describe the immunology that is specific to this disease. This is the gap that our work adresses.

The two potential outcomes of autoimmune myocarditis, resolution and chronic inflammation, are reminiscent of the “three Es” of immunoediting, where the interactions between a tumour and the immune system can lead to tumour elimination, equilibrium or escape [23]. This concept has been captured in mathematical models by systems of ODEs that describe the dynamic interactions between tumour and immune system components and have solutions that reveal the existence of multiple steady states. In these systems, each steady state represents one of the possible outcomes of tumour-immune interactions, i.e. one of the three Es [12].

Similarly, it is reasonable to expect a mathematical model of autoimmune myocarditis to have at least two non-negative steady states, one representing a healthy state, i.e. resolution, (green in Fig. 1) and one representing chronic inflammatory disease, with high numbers of immune cells and dead/damaged cardiomyocytes, or cardiac muscle cells (red in Fig. 1). A third non-negative steady state, a saddle point, separates their basins of attraction (orange in Fig. 1).We propose that the separatrix between the basins of attraction of the healthy and diseased steady states can be seen as a tolerance threshold in the immune system: any challenge to the immune system that stays below this threshold will be tolerated and will not lead to disease, i.e. the system stays in the basin of attraction of the healthy steady state (left panel in Fig. 1). However, if the threshold is crossed, autoimmune myocarditis develops. In a healthy individual, the body is able to tolerate small challenges to the immune system so the separatrix should not lie to close to the healthy steady state. Within this framework, we hypothesize that the impact of ICIs on the tolerance threshold reflects the increased risk of developing myocarditis when a cancer patient is treated with ICIs [4, 5]. Thus, the presence of ICIs should move the separatrix closer to the healthy steady state, and an immune challenge that could be controlled before treatment with ICIs, now leads to disease. This is shown in the left panel in Fig. 1 where the trajectories that approached the healthy steady state before (right panel), now lie above the lowered tolerance threshold and thus approach the diseased steady state instead. This hypothesis for the behavioural characteristics of a model of autoimmune myocarditis is tested in this paper.

**Figure 1:**
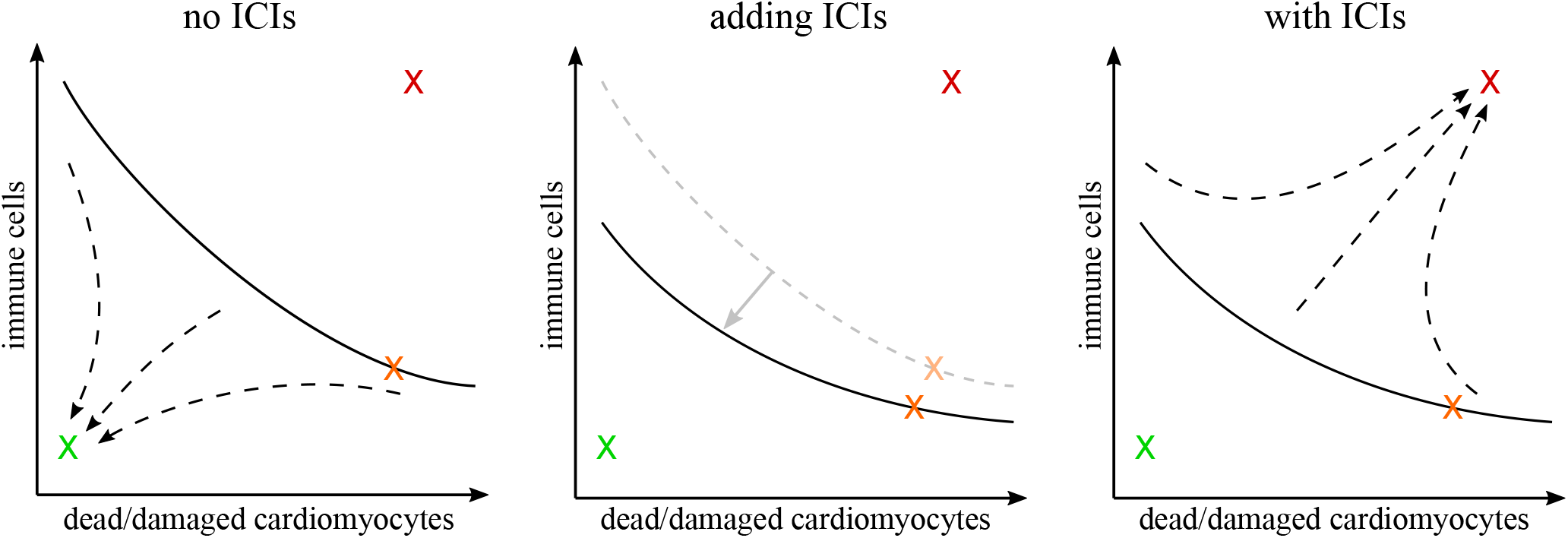
Illustration of the steady state and stability concepts related to the hypothesized mechanism of autoimmune myocarditis and the role of ICIs therein. The green cross indicates a healthy steady state with low cardiac damage and low numbers of immune cells. The red cross indicates a diseased steady state with high cardiac damage and high numbers of immune cells. The orange cross indicates a saddle point. The dashed arrows show trajectories, starting at the same initial condition in both the left and right panels. By adding ICIs (middle panel), the separatrix is moved closer to the healthy steady state, lowering the threshold for an immune response to develop into myocarditis.

### 1.1. Aims and outline

We propose a mathematical model of autoimmune myocarditis based on the known biological pathways underlying the disease, and explore whether such a model can produce the behaviours postulated in Fig. 1. The cell types included in this model represent a minimal set required to describe the mechanisms underlying the development and progression of autoimmune myocarditis. In addition to increasing our understanding of the immunology underlying autoimmune myocarditis, this model takes a first step towards designing patient-specific treatment regimes that balance the benefit of treating cancer with the risk of developing myocarditis.

Section 2 describes the background immunology needed to develop the model. Section 3 presents the model, the steady states and the parameterisation. In Section 4 we show that the presence of ICIs in the system increases the likelihood of a patient developing autoimmune myocarditis. Lastly, in Section 5 we explore how the administration of ICIs might affect the risk of autoimmune myocarditis for different patients by investigating model behaviour in different parameter regimes. We conclude in Section 6 with a brief discussion and proposal for avenues for future research.

## 2. Immunology

Here we provide a brief summary of the immunology of autoimmune myocarditis and the mechanisms of action of nivolumab and ipilimumab, two ICIs commonly used in the treatment of cancer. For more detail, the interested reader is referred to the recent review paper [24].

The damage to cardiomyocytes, cardiac muscle cells, that leads to the development of myocarditis can occur spontaneously or it can be triggered by viral infection, cross-reacting T cells, molecular mimicry, or drugs like ICIs, either acting alone or in combination [25]. When cardiomyocytes are damaged or dying their intracellular content can be released. One of the intracellular proteins found in cardiomyocytes is myosin heavy-chain alpha (MyHC*α*), which is unique to the myocardium and is believed to be one of the major antigens targeted in autoimmune myocarditis [6]. It is a cryptic antigen, meaning the body does not recognize it as “self” and produces an immune response against it [6].

Independent of what triggered the inflammation, sustained, chronic inflammation driven by the release of self-antigens leads to tissue remodeling and fibrosis, which in turn leads to cardiac dysfunction. This is known as dilated cardiomyopathy, the most severe consequence of myocarditis. Fibrosis is an irreversible process, and ultimately, the cardiac dysfunction caused by it will result in heart failure [9]. It is thus key that inflammation is resolved as quickly as possible in order to avoid this fatal outcome.

The immune response can be roughly divided into a fast, non-specific innate immune response and a slowly developing, antigen-specific adaptive immune response [26]. We briefly describe both below.

### 2.1. Innate immune response

The innate immune response relies on tissue-resident sentinel cells, such as macrophages and fibroblasts, and their signaling to recruit other immune cells, such as polymorphonuclear leukocytes (PMNs) [27]. Macrophages are present in high numbers at the inflammatory site in autoimmune myocarditis [28]. Their function depends on their phenotype, which lies on a spectrum between a pro-inflammatory extreme, M1, and an anti-inflammatory extreme, M2. Macrophages maintain high phenotypic plasticity and can change their behaviour in response to environmental cues. M1 macrophages specialise in antigen presentation, phagocytosis of pathogens and dead cells, and stimulate the T cell-mediated immune response, which is further discussed below. M2 macrophages secrete anti-inflammatory cytokines, stimulate wound healing and immune tolerance, and modulate the T helper (Th)2 cell pathway [9, 26]. The balance between these two phenotypes needs to be tightly regulated to ensure an effective immune response [29].

PMNs are a group of immune cells that include eosinophils, neutrophils, basophils and mast cells [26]. Eosinophils and neutrophils in particular have been implicated in the pathology of autoimmune myocarditis [30, 31]. Both of these immune cell types can respond to antigens by degranulating, a process in which they release toxic compounds into the extracellular space [30, 32]. Neutrophils further attack pathogens via phagocytosis or by trapping them in neutrophil extracellular traps [33, 34]. Eosinophils play an additional role in regulating the immune response through cytokine secretion [35].

Lastly, fibroblasts are tissue-resident stromal cells that play an important role in maintaining the structural integrity of the tissue. They are essential for wound healing and the development of scar tissue, which they achieve through remodeling of the extracellular matrix [26]. Heart tissue has little-to-no regenerative capabilities so any damaged tissue is replaced by scars, impeding heart function [26, 36]. Activation of fibroblasts early on in inflammation can limit the severity of autoimmune myocarditis. If, however, fibroblasts are active for too long, they can cause pathological scarring and tissue fibrosis, which impedes heart muscle function and eventually leads to heart failure [10].

The innate immune response is essential in the first few hours to days after tissue injury or infection [26]. It is characterized by high levels of immune-related proteins, PMNs, and edema fluid at the site of inflammation. If the injury or infection cannot be cleared by the innate immune response, the adaptive immune response, driven by antigen-specific T cells, takes over. We summarise this pathway in the next section.

### 2.2. Adaptive immune response

Although the adaptive immune response relies on both B and T cells, we focus here on CD4+ T cells as these have been identified as the drivers of the development and progression of autoimmune myocarditis [9].

#### 2.2.1. Antigen presenting cells

Cardiac antigens such as MyHC*α* that have been released due to cardiomyocyte damage are picked up by antigen presenting cells (APCs). Cells that can function as APCs include macrophages, monocytes and dendritic cells (DCs) [37]. They can be tissue-resident, as is the case for macrophages, or circulate through the body, as is the case for DCs [38]. APCs ingest pathogens, cell debris and other antigens, break them down and present fragments of them on major histocompatibility complexes on their cell surface [26, 39].

APCs drain to secondary lymph tissue or the spleen, where they come in contact with mature, naïve T cells [38, 40]. These T cells are ready to be activated by an APC that presents to them both the antigen that they recognize, as well as critical co-stimulatory proteins such as CD40, CD80, and CD86 [40]. If a T cell binds an antigen on an APC without simultaneously binding the required co-stimulatory proteins, it goes into anergy, a hyporesponsive state in which the T cell is unable to proliferate or produce cytokines [39, 41]. Because the CD4+ T cell-mediated pathway is central to ICI-induced autoimmune myocarditis, we focus here on CD4+ T cells and use “CD4+ T cells” and “T cells” interchangeably.

#### 2.2.2. CD4+ T cells

After activation by an APC, CD4+ T cells go through a process of clonal expansion, during which they divide approximately seven times [42]. The daughter cells produced as part of this process differentiate into a number of phenotypes, most importantly Th and T regulatory (Treg) cells. The phenotype that the T cells assume depends largely on soluble mediators produced by APCs and innate immune cells [39, 41, 42, 43].

There are a number of Th cell subtypes, all of which have a predominantly pro-inflammatory function. Th1, Th2 and Th17 cells play the most important role in autoimmune myocarditis. Together, these subtypes stimulate the recruitment and activation of macrophages and PMNs, most importantly eosinophils and neutrophils, influence the differentiation of newly activated CD4+ T cells, inhibit each others activity levels, stimulate DC maturation, and damage inflammed tissue, amongst other functions [26, 39, 44, 45].

In contrast, Treg cells are anti-inflammatory T cells that inhibit the proliferation, survival and cytokine secretion of DCs, T cells and innate immune cells, such as PMNs [46, 47]. They use a number of different pathways, including cell-cell contact, cytokines and resource competition to control the immune response by influencing the activity of Th cells, DCs, and innate immune cells [48, 49]. APCs and T cells thus form an immune pathway targeted at a specific antigen.

Taken together, the innate and adaptive immune responses, and the many links between them, form the basics of the immunology underpinning the development and progression of autoimmune myocarditis. For clarity, much detail has been omitted in this summary, but even so the immense complexity of the underlying mechanisms of autoimmune myocarditis is clear. Fig. 2 provides a graphical overview of the immunology discussed above.

**Figure 2:**
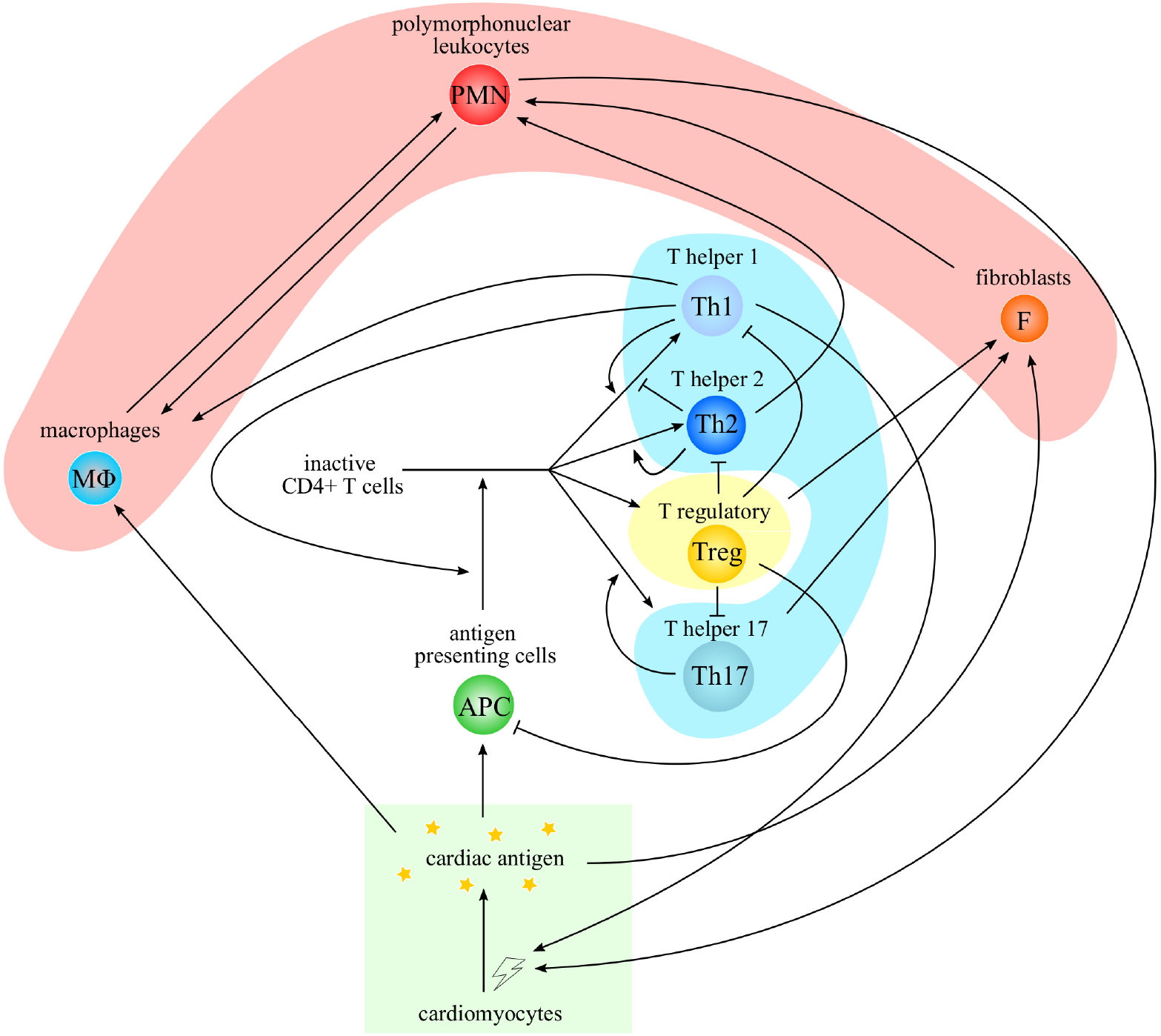
An overview of the immunology underpinning the development and progression of autoimmune myocarditis. Arrowheads indicate stimulation, barheads indicate inhibition. Shaded areas represent the lumped variables in the mathematical model.

### 2.3. Immune checkpoint inhibitors

The motivation behind ICIs as a cancer therapy is to leverage the powerful immune response against malignancies like tumours. To do this, inhibitory pathways affecting the activation and inhibition of immune cells like CD4+ T cells, are blocked using therapeutic antibodies. Immune-related adverse events arise as side-effects of ICIs when the disinhibited immune response targets non-malignant tissues. Cardiotoxicity, when the immune response is misguided to the heart, is an uncommon irAE, but one with an often sudden onset and quick escalation [7]. The time to onset of symptoms of ICI-induced autoimmune myocarditis varies widely between patients, with some showing symptoms as early as three days after infusion and others taking over a year to display symptoms. The median time to onset is 30–65 days, which corresponds to between one and three doses of ICIs on current treatment schedules [50].

Ipilimumab is one ICI with autoimmune myocarditis as a potential side effect [51, 52]. It is an antibody against cytotoxic T-lymphocyte-associated protein 4 (CTLA4). CTLA4 is found on the surface of activated T cells. On Treg cells in particular it is constitutively expressed [46, 53]. CTLA4 binds CD80 and CD86, the co-stimulatory proteins on the surface of DCs, with higher affinity than CD28, which is present on the surface of inactive T cells [43]. Ipilimumab has two main mechanisms of action. Crystallographic evidence shows that the binding site of ipilimumab on CTLA4 overlaps with the binding site of CD80 and CD86 [54, 55]. Binding of ipilimumab to CTLA4 thus creates a steric block so that CTLA4 can no longer bind CD80 and CD86. Therefore, in the presence of ipilimumab, CD28 has less competition for binding sites on APCs, leading to more positive stimulation of T cells by APCs. Blocking CTLA4 with ipilimumab also leads to the depletion of Treg cells [54, 56]. Both of these mechanisms, the blocking of inhibitions of T cell activation and the depletion of Treg cells, lead to an increase in T cell-mediated inflammation, both in the tumour environment and other tissues.

Another ICI with autoimmune myocarditis as a known side effect is nivolumab, which has a different mechanism of action [51]. Nivolumab is an antibody against a protein called programmed cell death protein-1 (PD-1). PD-1 is carried by Th cells on their cell surface [57]. One of its ligands, PD-L1, is found on the surface of peripheral cells, including cardiomyocytes, where it is expressed in response to pro-inflammatory cytokines. A second ligand, PD-L2, is expressed on DCs in response to antigen uptake. Its binding by PD-1 protects the DC against the cytotoxicity of activated T cells and prevents their overactivation [57, 58]. When PD-1 binds one of its ligands, the T cell is inhibited and anergy is induced [53]. PD-1-blocking therapy leads to an increase in the expression of proteins for proliferation, activation and effector functions in CD4+ T cells, and thus increases both anti-tumour T cell activity and the probability of inflammation in other organs [59].

Finally, we note that autoimmune myocarditis is often caused by a combination of triggers. Although ICIs increase the risk of developing this disease by increasing T cell activation and decreasing inhibition, they are generally not sufficient to trigger it alone. Often cardiac antigen-specific T cells are in circulation, either due to low negative selection in the thymus or because tumour and gut bacteria-specific T cells are cross-reactive [53, 60]. These cells act as secondary trigger for the development of autoimmune myocarditis.

To better direct future research to avenues that are most promising, we need to determine which immunological processes are essential for the development and progression of autoimmune myocarditis, which are of secondary importance, and what roles ICIs play in driving the disease. To aid in making these determinations, we have developed a mathematical model of this disease, which we present in the next section. This model strikes a balance between simplicity, to allow analytical methods to be applied in its study, incorporating cell types determined to be essential by experimental research, and biological realism, to preserve disease characteristics observed in patients.

## 3. The model

Here, we introduce the model for autoimmune myocarditis and the effects of ICIs. Our guiding aim is to capture the salient immunological mechanisms and preserve the characteristics of the disease observed in patients while keeping the model tractable. The relative simplicity of ODE models combined with their ability to capture complex, highly nonlinear immunological mechanisms, makes them a suitable starting point for modelling this disease.

Fig. 2 shows the complexity of the key immunological interactions. However, including all these cell types and interactions would lead to a highly complex model and preclude a number of approaches that provide insight into the behaviour of the system, such as the derivation of analytical expressions for the steady states and their linear stability. Instead, we group together immune cell types that have similar effects on the cycle of autoimmunity. For example, Th cells, shaded in blue in Fig. 2, all have a similar function and can thus be grouped together. Similarly, the innate immune cell types, i.e. macrophages, fibroblasts and PMNs shaded in red, are all activated by T cells or directly by damage to the myocardium and ultimately lead to activated PMNs attacking cardiomyocytes, and thus we also group these cell types together. Although this inevitably leads to the ignoring of certain immunological details, our goal is to build a minimal model that includes the immunological pathways required to capture the development of autoimmune myocarditis.

Our model consists of a system of four nonlinear ODEs, each describing one class of cell types: the number of damaged/dead cardiomyocytes (*C*), an indicator of the extent of cardiac damage and disease progression; the number of innate immune cells (*I*); the number of pathogenic CD4+ T cells (*P*); and the number of regulatory T cells (*R*). The ICIs nivolumab and ipilimumab are represented in the system by the unitless variables *D*_*N*_ and *D*_*I*_, respectively. Fig. 3 shows a graphical representation of the model. Table 1 lists the parameters with descriptions and units. The variable *t* denotes the time in units of days.

**Table 1:**
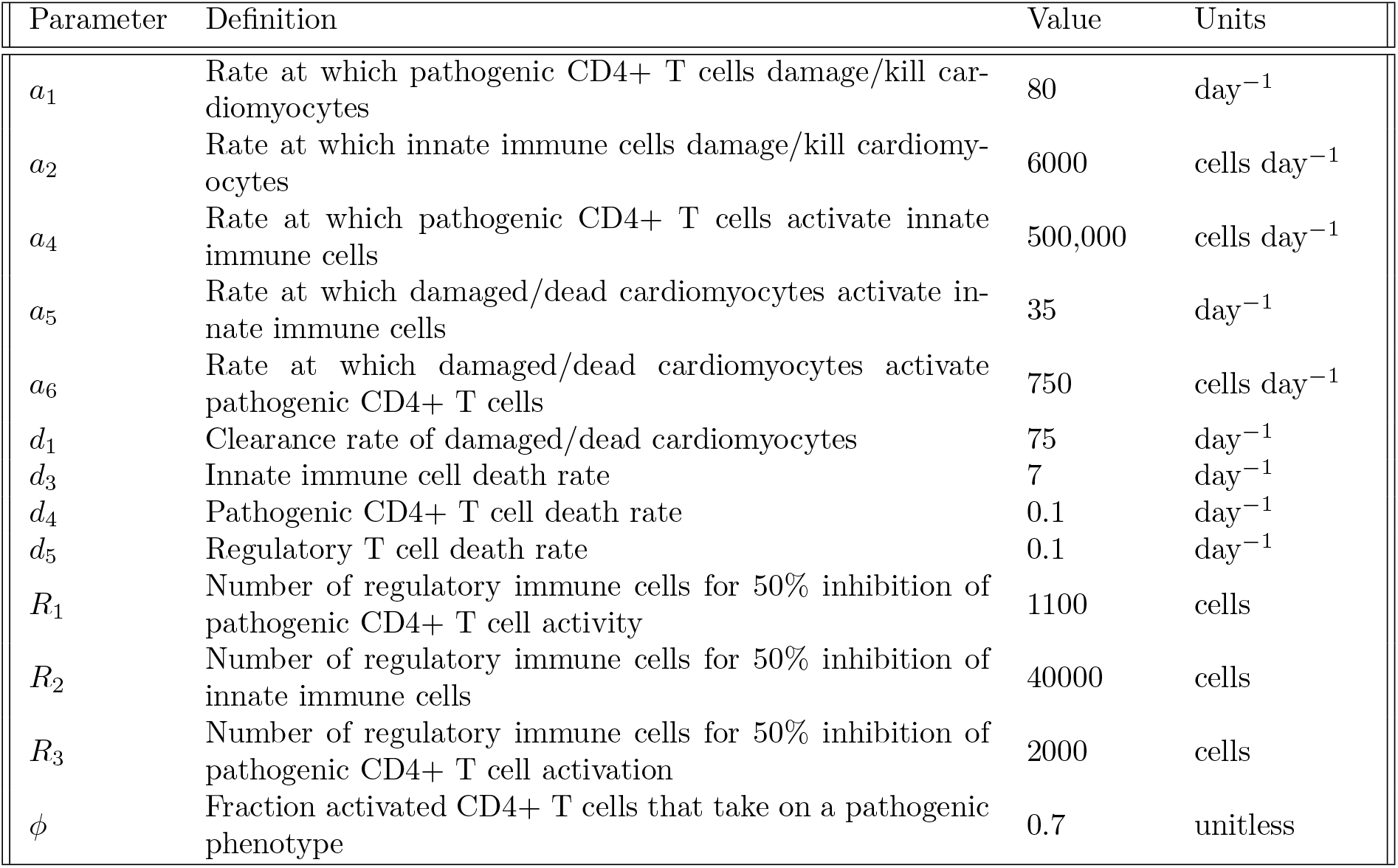
Definitions, values and units of the 13 model parameters.

**Figure 3:**
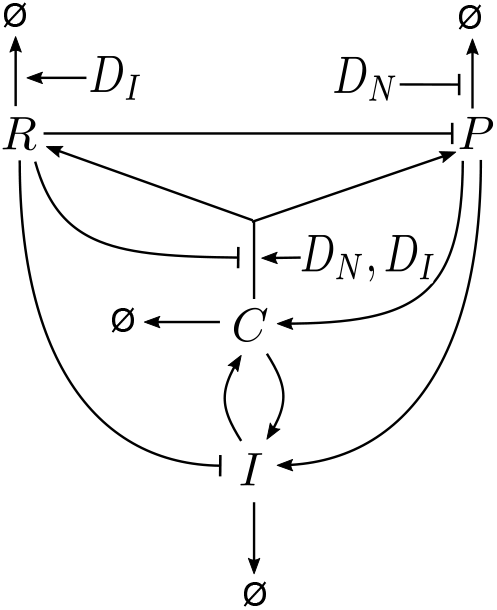
Graphical representation of the mathematical model of autoimmune myocarditis, indicating the effects of nivolumab (*D*_*N*_) and ipilimumab (*D*_*I*_) in the network of interactions. The cell types included are dead/damaged cardiomyocytes (*C*), innate immune cells (*I*), pathogenic CD4+ T cells (*P*) and regulatory T cells (*R*). Arrow-heads indicate stimulation, bar-heads indicate inhibition, Ø indicate cell death and/or clearance.

### 3.1. The mathematical model

The dynamics of dead/damaged cardiomyocyte numbers are governed by

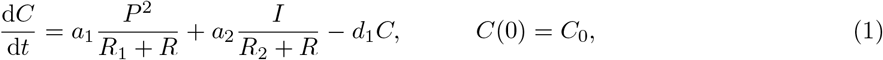

where *a*_1_, *a*_2_, *R*_1_, *R*_2_ and *d*_1_ are positive parameters, and *C*_0_ is the nonnegative initial number of damaged cardiomyocytes. The first term on the right-hand side (RHS) of Eq. (1) describes the nonlinear effects of damage to the myocardium due to the presence of pathogenic CD4+ T cells. The denominator *R*_1_ + *R* represents the inhibition of this process by Treg cells, where *R*_1_ is the number of regulatory immune cells required for 50% inhibition of pathogenic CD4+ T cell activity. The second term on the RHS of Eq. (1) describes myocardial damage due to the presence of innate immune cells [9, 31, 45, 61]. This process is inhibited by Treg cells, hence the denominator, where *R*_2_ is the number of regulatory immune cells required for 50% inhibition of innate immune cells. Lastly, the third term on the RHS of Eq. (1) describes the rapid clearance of dead cardiomyocytes [62, 63]

The number of innate immune cells evolves over time according to the ODE

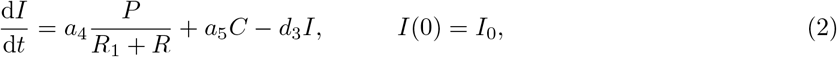

where *a*_4_, *a*_5_ and *d*_3_ are positive parameters, and *I*_0_ is the nonnegative initial number of innate immune cells. The first term on the RHS of Eq. (2) describes the stimulation of innate immune cells by pathogenic CD4+ T cells [45, 64]. Again, pathogenic CD4+ T cells are inhibited by Treg cells. The second term on the RHS of Eq. (2) describes the recruitment of innate immune cells by dead/damaged cardiomyocytes through the innate immune pathway [26, 65, 66]. The last term describes the death or deactivation of innate immune cells [67, 68].

In this model, the pathogenic CD4+ T cell population is assumed to consist primarily of Th cells. The dynamics of pathogenic CD4+ T cells is governed by the ODE

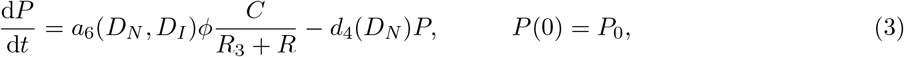

where *ϕ* and *R*_3_ are positive parameters, *P*_0_ is the nonnegative initial number of pathogenic T cells, *a*_6_(*D*_*N*_, *D*_*I*_) is a monotonically increasing function of both nivolumab and ipilimumab doses, and *d*_4_(*D*_*N*_) is a monotonically decreasing function of nivolumab dose. In an ICI-free environment, *a*_6_(0, 0) > 0 is the activation rate of T cells due to cardiac damage and *d*_4_(0) > 0 is the deactivation/death rate of pathogenic CD4+ T cells. The first term on the RHS describes the activation of inactive T cells by dead/damaged cardiomyocytes and their subsequent clonal expansion [26, 69]. A fraction 0 < *ϕ* < 1 of activated T cells then adopt a pathogenic phenotype [6, 26, 70]. The processes of activation and differentiation are inhibited by Treg cells, hence the denominator *R*_3_ +*R* where *R*_3_ is the number of regulatory immune cells required for 50% inhibition of pathogenic CD4+ T cell activation [26, 46, 48, 61]. The second term on the RHS of Eq. (3) denotes the deactivation or death of pathogenic CD4+ T cells [7, 9, 57, 58, 71, 72]. As discussed above, nivolumab prevents T cell inhibition in the myocardium and both nivolumab and ipilimumab stimulate T cell activation. These mechanisms of action are reflected in the model by the effects of *D*_*N*_ and *D*_*I*_ on the parameters *a*_6_ and *d*_4_.

The number of Treg cells evolves according to the ODE

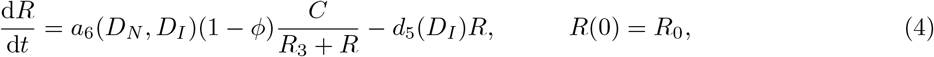

where *ϕ* and *R*_3_ are positive parameters, *R*_0_ is the nonnegative initial number of Treg cells, and *d*_5_(*D*_*I*_) is a monotonically increasing function of ipilimumab dose with *d*_5_(0) > 0 the death rate of Treg cells in an ICI-free environment. The activation and differentiation of Treg cells happens according to the same processes as described for pathogenic CD4+ T cells, the only difference being that a fraction (1 − *ϕ*) of the clonally expanded T cells adopt a regulatory phenotype. This is reflected in the first term on the RHS of Eq. (4). The second term on the RHS of Eq. (4) describes the death or deactivation of Treg cells. Ipilimumab depletes Treg cells and this is reflected in the model by the fact that *d*_5_ is a function of *D*_*I*_, which increases the death rate of Treg cells.

### 3.2. Steady states and linear stability

The model described in Section 3.1 was built based on the immunology underlying the progression of autoimmune myocarditis. To assess whether this model can express the behaviour that we hypothesized to be characteristic of a model for this disease (see Section 1. and Fig. 1), we now seek an analytical expression for the steady states of the system. As discussed in Fig. 1, a model of autoimmune myocarditis should allow three non-negative steady states: one stable steady state with low counts of inflammatory cells and dead/damaged cardiomyocytes; one stable steady state with high counts of inflammatory cells and dead/damaged cardiomyocytes; and one saddle point with intermediate cell counts.

The origin is a steady state of the system (1)-(4). To determine the non-zero steady states of the model, we set the RHSs of Eq. (1)-(4) equal to zero, and use algebraic manipulation to eliminate *C, I* and *R*, and establish a polynomial equation satisfied by the non-zero steady states in terms of *P*. The non-zero steady state numbers of pathogenic CD4+ T cells, *P*_∗_, satisfy the cubic equation

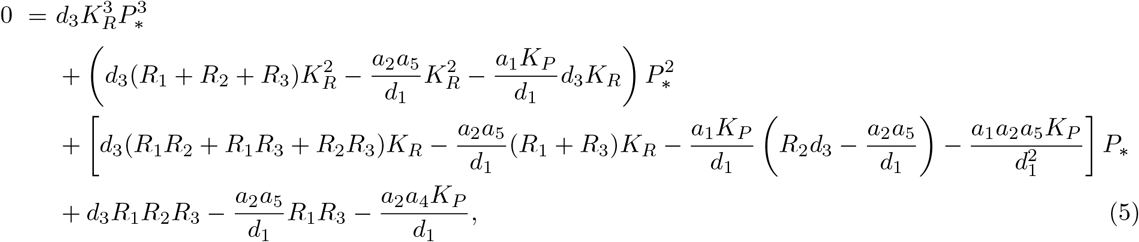

where 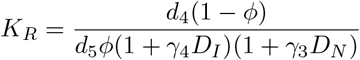 and 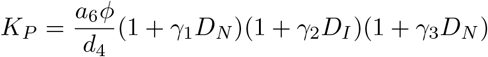.

The steady state cell counts of the remaining variables are then given by

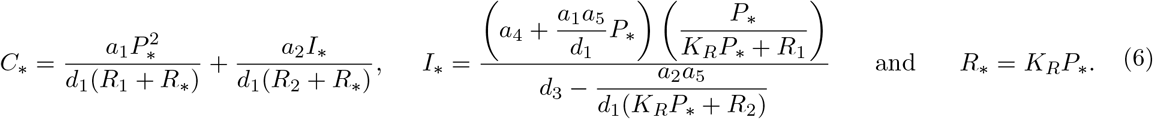

The linear stability of the non-negative steady states is determined by the eigenvalues of the Jacobian matrix

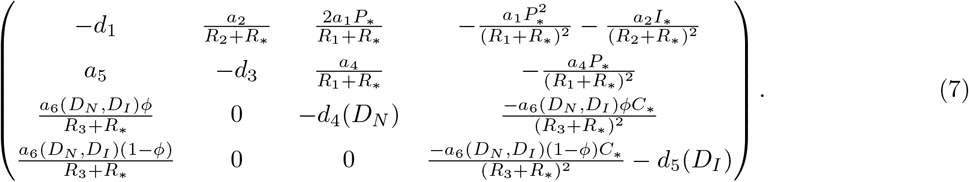

### 3.3. Parameters

Our overarching aim is to develop a model capable of recapitulating the increased risk of developing autoimmune myocarditis upon application of ICIs. The mathematical model has 13 parameters (see Table 1). The grouping of different types of immune cells into classes and the consolidation of many steps in the immunological process into single terms in the model means the model is simple enough to admit some analytical approaches, but it makes it difficult to estimate most parameter values directly from the literature. Instead, we seek parameter values that give rise to the hypothesized number and types steady states required for the model behaviour illustrated in Fig. 1, in an ICI-free context (*D*_*N*_ = *D*_*I*_ = 0). The values of parameters *d*_4_ and *d*_5_ can be obtained from literature [73]. For the remaining 11 parameters, we find suitable ranges based on a combination of literature insights and constraints that give rise to the behaviour depicted in Fig. 1. Recall, we proposed that a model of autoimmune myocarditis should have two non-negative, stable steady states and one saddle point, whose position in phase space is influenced by the presence of ICIs. In this section, we explore whether, and for what parameter values, the model can display these characteristics. We seek parameter sets that satisfy the following criteria (summarised in Table 2):

**Table 2:** Requirements for establishing parameter values.

| Requirement | Reference |
| --- | --- |
| Three nonnegative steady states, two stable and one saddle. | Fig. 1 |
| Number of damaged/dead cardiomyocytes, pathogenic CD4+ T cells and Treg cells at the saddle point must be between $10^2$ and $10^4$. | [42, 74] |
| Number of damaged/dead cardiomyocytes should never exceed $10^6$. | [75] |
| Numbers of T cells peak around seven days. | [13] |
| All terms must have significant impact on model output. | Section 2 |
| $d_4 = d_5 = 0.1 \text{ day}^{-1}$. | [73] |

1. The parameter set must produce three nonnegative steady states. Two must be stable to small perturbations, and one must be a saddle point so that the behaviour of the model captures the ideas in Fig. 1. The origin corresponds to the “healthy” state, with no inflammatory cells. This steady state must be stable. At the origin where [*C*_∗_, *I*_∗_, *P*_∗_, *R*_∗_] = [0, 0, 0, 0], the Jacobian matrix is

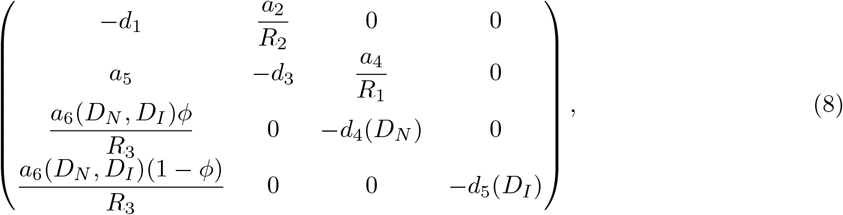

from which we derive the following condition for stability:

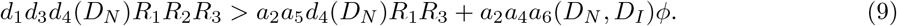
2. The saddle point cell counts of damaged cardiomyocytes, innate immune cells and pathogenic CD4+ T cells should be between 10^2^ and 10^4^ cells, to ensure that a moderately sized perturbation in CD4+ T cells is needed to launch an immune response. Although it is unknown how many immune cells or dead/damaged cardiomyocytes are needed to trigger an immune response, it is known that approximately 3000 DCs are required for a 50% chance of launching an immune response, and we assume that the threshold is at a similar cell count for the other cell types [42, 74].
3. The maximum number of damaged cardiomyocytes at any time point must be below 10^6^. The heart consists of approximately 2 − 3 *×* 10^9^ cardiomyocytes in total. To be biologically realistic, the model should not predict the entire heart to be destroyed within a matter of days [75].
4. The number of T cells should peak around seven days after an immune challenge [13].
5. All terms in the model must have a significant impact on model outcomes. Because the model was constructed to include those interactions which, based on the existing literature, are important for the development of autoimmune myocarditis, all these interactions should have a significant impact on the dynamics.

We obtain estimates for *d*_4_ and *d*_5_ from Robertson-Tessi et al. [73], setting *d*_4_ = *d*_5_ = 0.1 day ^−1^. We implement a parameter search to establish reasonable values for the 11 remaining parameters. All algorithms, here and in the following sections, are implemented and all simulations run in MATLAB R2020b (published by Mathworks, Natick, MA). Unless noted otherwise, we use the ode23s solver with default tolerances. A stiff solver is required because of the rapid changes in cell counts over time, and we note that the ODE solutions converge for the default tolerances.

To find parameter sets that satisfy criteria one and two, we solve Eq. (5). By Descartes Rule of Signs, the number of sign changes in the coefficients of the polynomial in Eq. (5) gives the maximum number of positive solutions for *P*_∗_. Eq. (5) can be solved using the roots() function in MATLAB to obtain the nonzero steady state cell counts of *P*_∗_. After solving for the steady state cell counts of the other variables, we then select those steady states with suitable positive cell counts for all variables and the desired linear stability.

To further narrow down the number of feasible parameter sets to those satisfying criteria three and four, we simulate the model using the remaining parameter sets using the initial condition [*C*_0_, *I*_0_, *P*_0_, *R*_0_] = [0, 0, 10^4^, 0] cells. The healthy steady state is at the origin. Because active, cardiac-specific pathogenic CD4+ T cells are a known (partial) trigger of autoimmune myocarditis, setting an initial condition of *P*_0_ = 10^4^ is employed as a means to perturb the system away from a healthy state as this number of pathogenic T cells will ensure the development of autoimmune myocarditis, as required for testing criteria three and four [53, 60]. From these simulations, we extract the maximum number of damaged/dead cardiomyocytes in the system at any point and discard those parameter sets for which this cell count is too high. Next, we select parameter sets for which the number of T cells in the system peaks as close to seven days as possible. The remaining parameter sets thus satisfy criteria three and four. We then select one set of parameter values from the many that satisfy the criteria listed above to make progress on modelling autoimmune myocarditis. The parameter values in this “default parameter” set are listed in Table 1.

To confirm that the default parameter set satisfies criterion five, we apply a parameter sensitivity analysis on an ICI-free model, i.e. setting *D*_*N*_ = 0 and *D*_*I*_ = 0, using the implementation of Latin Hypercube Sampling Partial Rank Correlation Coefficent (LHS-PRCC) made available by Marino et al. [76]. This metric tells us how much a particular output of the model is affected by an increase or decrease in a certain parameter value, and whether or not this impact is significant compared to a dummy parameter. We choose this method because the computational costs are relatively low compared to other methods, such as the extended Fourier Amplitude Sensitivity Test, for such high-dimensional parameter spaces.

We take the initial conditions for the parameter sensitivity analysis to be [*C*_0_, *I*_0_, *P*_0_, *R*_0_] = [0, 0, 10^4^, 0] cells, so the system is perturbed away from the healthy steady state and towards autoimmune myocarditis, and the effect of the parameters on the development of the disease can be assessed. The impact that the parameters have on each of the variables is measured after 60 days, a relevant timescale for the development and pathology of autoimmune myocarditis, as discussed in Section 2.3. We vary each of the parameter values over two orders of magnitude around their default values. The only exception to this is *ϕ* which is constrained between zero and one. A new parameter set is generated for each run, where the parameter values are independently varied around their default value. Approximately 6,000 runs were required for estimates of the correlation between the parameter values and model outcome to converge for all parameters. Results in Fig. 4 show that all variables are sensitive to the values of all parameters, and that this parameter set thus satisfies criterion five.

**Figure 4:**
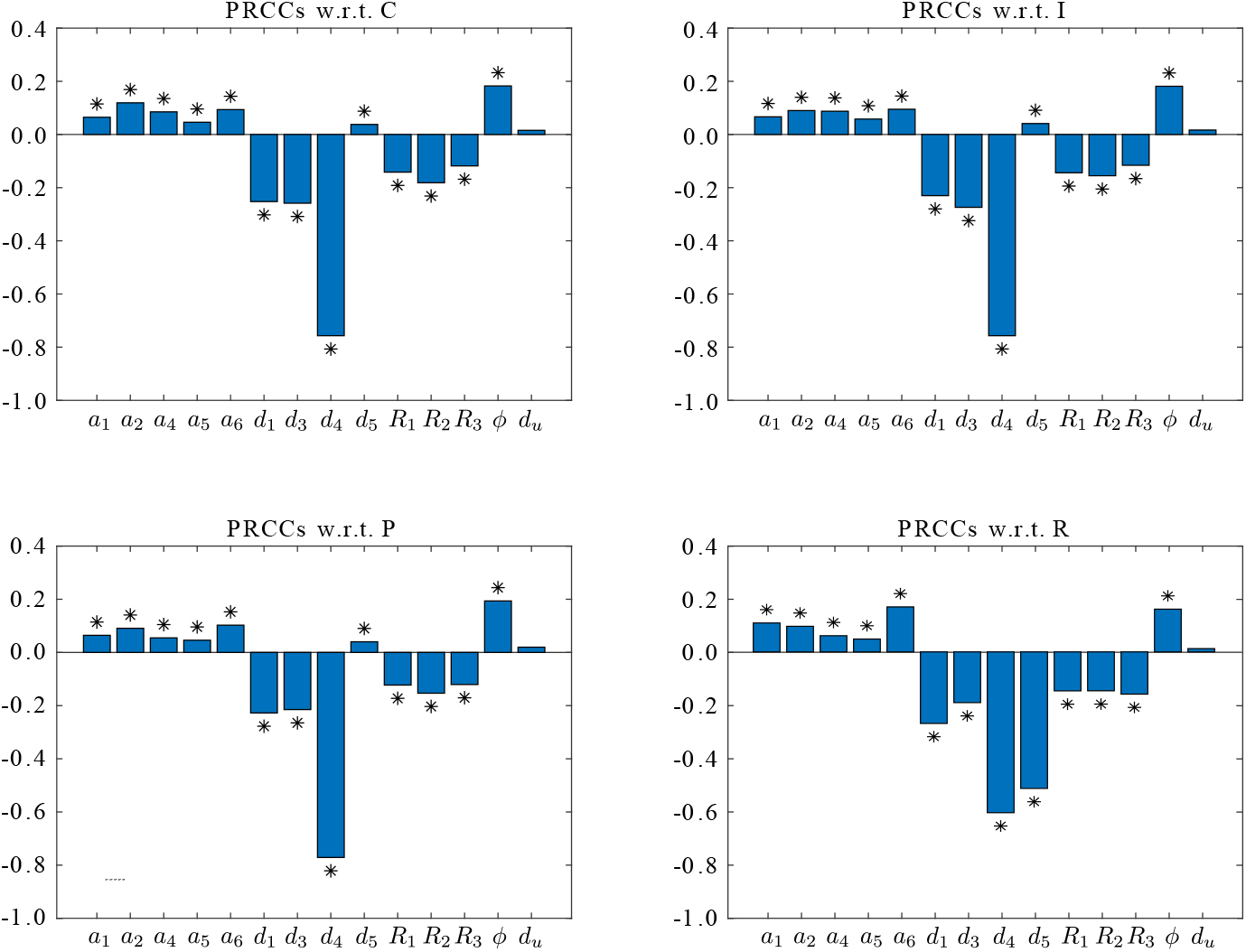
Parameter sensitivity analysis with baseline parameter values as given in Table 1. In each plot, *d*_*u*_ denotes the dummy parameter. Results obtained for initial conditions [*C*_0_, *I*_0_, *P*_0_, *R*_0_] = [0, 0, 10^4^, 0] cells, at *t* = 60 days and for 6,000 runs. Asterisk indicates a PRCC value that is significantly different from the dummy parameter, *p* ≤ 0.01, and the parameter thus has a significant impact on the output variable.

### 3.4. Model simulation with default parameter set

We expect model behaviour to change significantly depending on the magnitude of the perturbation away from the healthy steady state. For small perturbations, the system should be able to recover to the healthy steady state, but for perturbations above a certain threshold the system should tend to the diseased steady state. To explore this, we simulate the model with the default parameter set, setting all initial conditions to zero, except for *P*_0_ which is varied to simulate different immune perturbations away from the healthy steady state at the origin. The time traces in Fig. 5 show that between 250 and 500 pathogenic CD4+ T cells are required to trigger an immune response, which falls within the range of 100-10,000 T cells estimated based on existing literature [42, 74]. When autoimmune myocarditis develops, the T cell response takes approximately seven days to launch, while the innate immune response develops almost instantaneously. These behavioural characteristics confirm that the model, simulated using the default parameter set, is capable of reproducing experimental observations [13, 26]. Thus, with this particular parameter set, the minimal set of immune cell types and pathways included in this model suffices to recapitulate the disease characteristics seen in patients, and to produce the behaviour required from a mathematical model of autoimmune myocarditis, as postulated in Fig. 1, in an ICI-free context.

**Figure 5:**
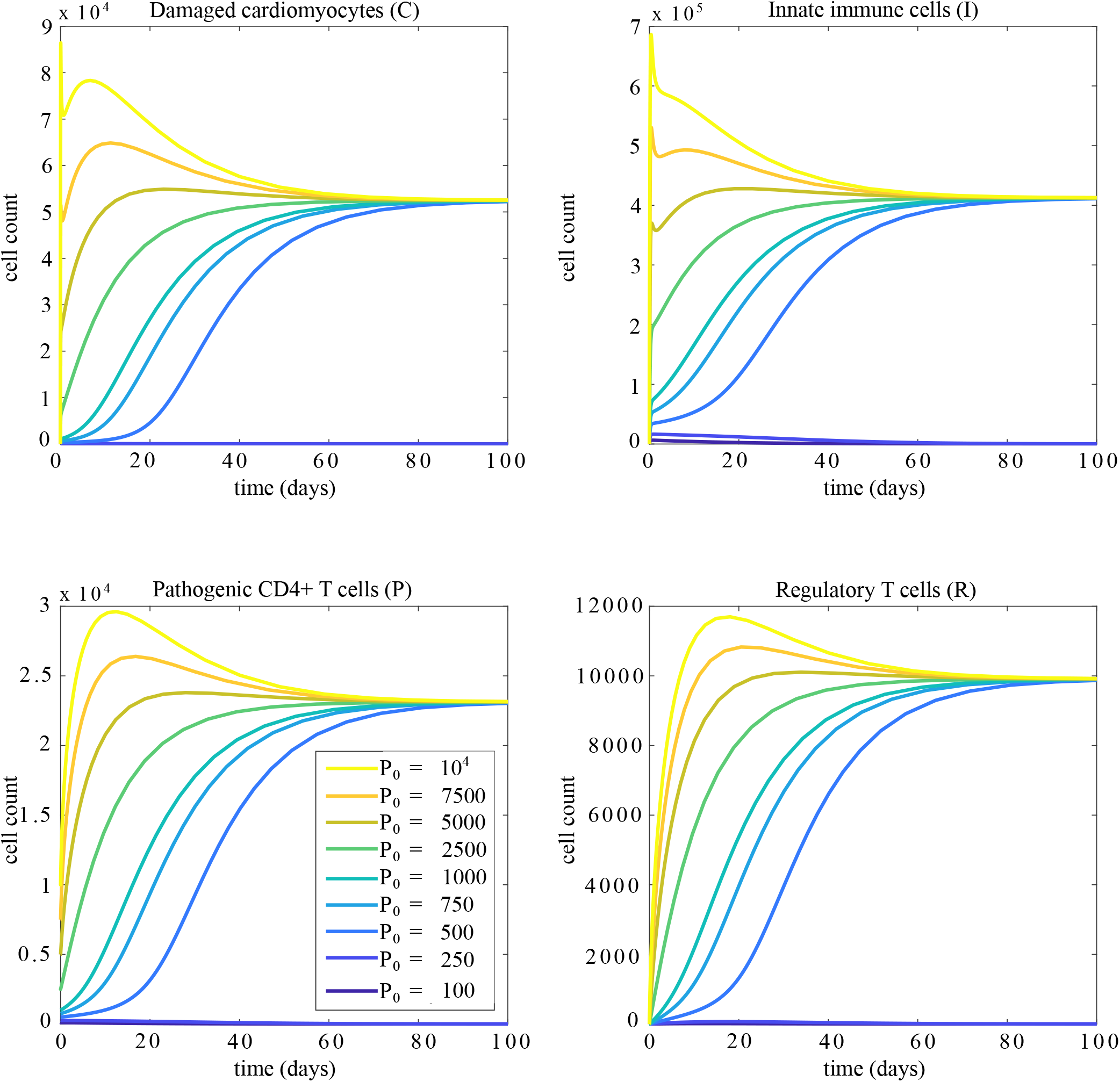
Time traces of the model variables for different initial conditions with default parameter values. The initial conditions are [*C*_0_, *I*_0_, *R*_0_] = [0, 0, 0] cells, with *P*_0_ varied as noted.

## 4. The effects of ICIs

In the previous section we established a mathematical model that is rooted in the experimental literature, and can replicate observations of the development of autoimmune myocarditis. We now turn to consider the effects that ICIs have on the system by exploring changes in the number and nature of the steady states as nivolumab and ipilimumab are administered.

Bifurcation structures provide insight into how the number of steady states and their linear stability change as a function of one or more model parameters. In the ICI-free model, the two stable steady states represent the two possible outcomes of autoimmune myocarditis: resolution and chronic inflammation. The changing values of *a*_6_(*D*_*N*_, *D*_*I*_), *d*_4_(*D*_*N*_) and *d*_5_(*D*_*I*_) as ICIs are administered affect the number and linear stability of the steady states, and thus how the system evolves after an immune challenge. As we are interested in how the steady states change qualitatively, we do not consider specific functional forms for *a*_6_(*D*_*N*_, *D*_*I*_), *d*_4_(*D*_*N*_) and *d*_5_(*D*_*I*_), but assume that the values of these parameters change monotonically as the doses of ipilimumab and nivolumab are increased.

### 4.1. Nivolumab

The left panel in Fig. 6 shows what happens to the number of steady states when nivolumab is applied to the system, changing the values of *a*_6_(*D*_*N*_, *D*_*I*_) and *d*_4_(*D*_*N*_). As previously discussed, nivolumab increases the rate of activation of T cells and decreases the rate of deactivation/death of pathogenic CD4+ T cells. Thus, as the dose of nivolumab is increased, the value of *a*_6_(*D*_*N*_, *D*_*I*_) increases and the value of *d*_4_(*D*_*N*_) decreases, moving the system from the default parameter set (starred in the left panel in Fig. 6) towards the bottom right-hand corner. If enough nivolumab is applied, the system crosses a bifurcation which engenders a switch from three steady states to only two steady states.

**Figure 6:**
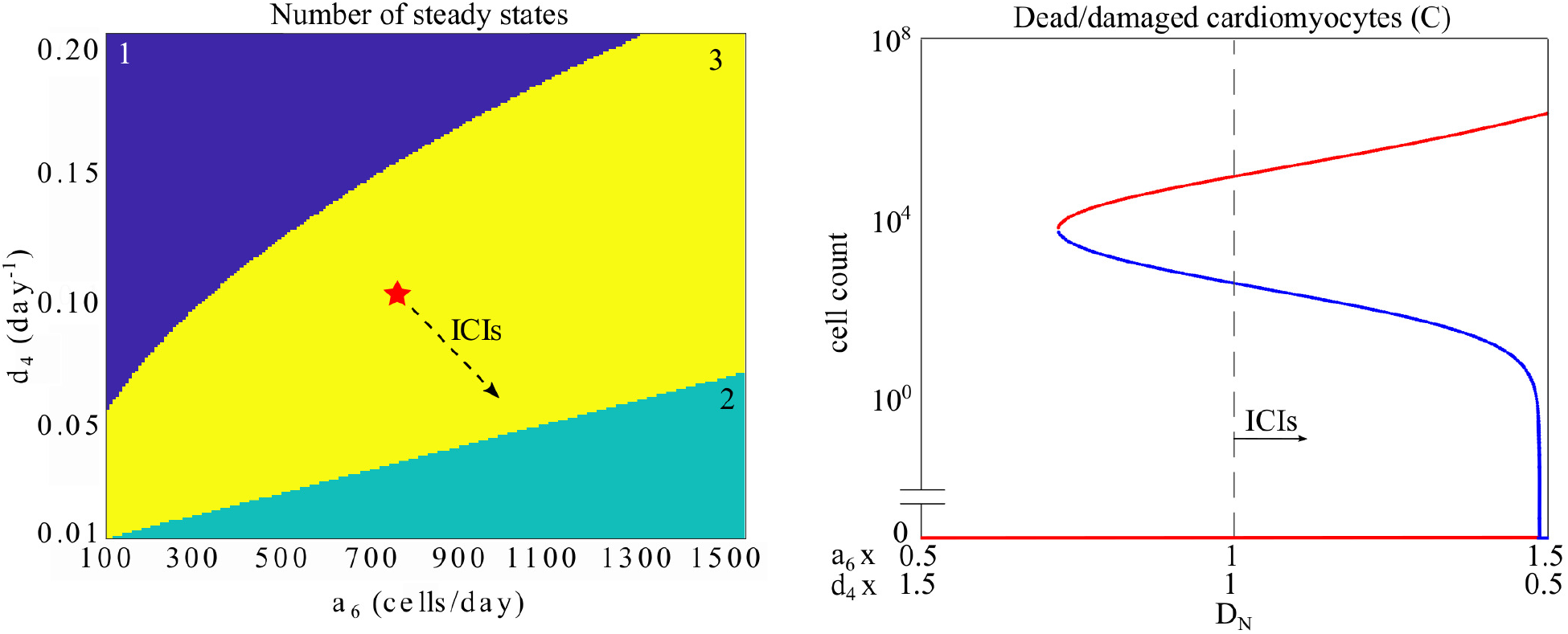
Left: Bifurcation diagram for varying values of *a*_6_(*D*_*N*_, *D*_*I*_) and *d*_4_(*D*_*N*_) as all other parameters are kept at default values. The yellow shading indicates regions of parameter space where the mode has three steady states. The green shading indicates regions with two steady states, and the purple shading indicates regions with one steady state. The red star indicates the default values of *a*_6_(*D*_*N*_, *D*_*I*_) and *d*_4_(*D*_*N*_). The dashed arrow indicates the general direction the system moves in as an increasing dose of nivolumab is applied. Right: A transcritical bifurcation occurs as the value of *a*_6_(*D*_*N*_, *D*_*I*_) is increased from its default value of 750 cells day^−1^ and the value of *d*_4_(*D*_*N*_) is decreased from its default value of 0.1 day^−1^. Default values are used for all other parameters. Red indicates a stable steady state, blue indicates a saddle point. The grey dashed line indicates the default values of *a*_6_(*D*_*N*_, *D*_*I*_) and *d*_4_(*D*_*N*_). The dashed arrow indicates the direction the system moves in when nivolumab is applied.

To explore the type of bifurcation that occurs as nivolumab is applied, the values of *a*_6_(*D*_*N*_, *D*_*I*_) and *d*_4_(*D*_*N*_) are varied and the steady states and their linear stability are tracked. It is assumed here that the effect of nivolumab on these parameters is identical in terms of the percent change in parameter value as a function of nivolumab dose. The right panel in Fig. 6 shows what happens to the stable steady states (red) and saddle point (blue) as the values of *a*_6_(*D*_*N*_, *D*_*I*_) and *d*_4_(*D*_*N*_) are varied between 50% and 150% of their default values. The dashed line in right panel in Fig. 6 indicates an ICI-free system, where *a*_6_(*D*_*N*_, *D*_*I*_) and *d*_4_(*D*_*N*_) are at their default values; here there are three steady states, two stable steady states and one saddle point, as established in previous sections. As nivolumab is applied in increasing amounts the cell count at the saddle point decreases, and the saddle point moves towards the origin. This means that fewer active immune cells are required to push the system past the separatrix, thus applying nivolumab lowers the threshold for the development of autoimmune myocarditis. As *a*_6_(*D*_*N*_, *D*_*I*_) nears 150% of its default value and *d*_4_(*D*_*N*_) nears 50% of its default value, the saddle point collides with the origin in a transcritical bifurcation, leaving the origin unstable. Past this bifurcation point, the system only has two steady states: a stable steady state with high immune cell counts and an unstable steady state at the origin. This means that any perturbation to the system away from the healthy steady state leads to the development of autoimmune myocarditis. Applying nivolumab to the system thus initially increases the likelihood of developing autoimmune myocarditis by lowering the threshold for immune perturbations to push the system across the separatrix until, for a high enough dose, any perturbation will lead to disease because the origin becomes unstable.

### 4.2. Ipilimumab

As the dose of ipilimumab is increased, both the activation rate of T cells, *a*_6_(*D*_*N*_, *D*_*I*_), and the death rate of Treg cells, *d*_5_(*D*_*I*_), increase. The left panel in Fig. 7 demonstrates that the system moves away from the default parameter set (starred) towards the top-right corner as more ipilimumab is applied. As with nivolumab, the system has three steady states for the default parameter set and crosses a bifurcation as an increasing dose of ipilimumab is applied, leaving the system with two steady states. In contrast to nivolumab, only one parameter, *a*_6_(*D*_*N*_, *D*_*I*_), controls the occurence of this bifurcation. For large values of *a*_6_(*D*_*N*_, *D*_*I*_), the number of steady states changes from three to two, but increasing only *d*_5_(*D*_*I*_) does not change the number of steady states.

**Figure 7:**
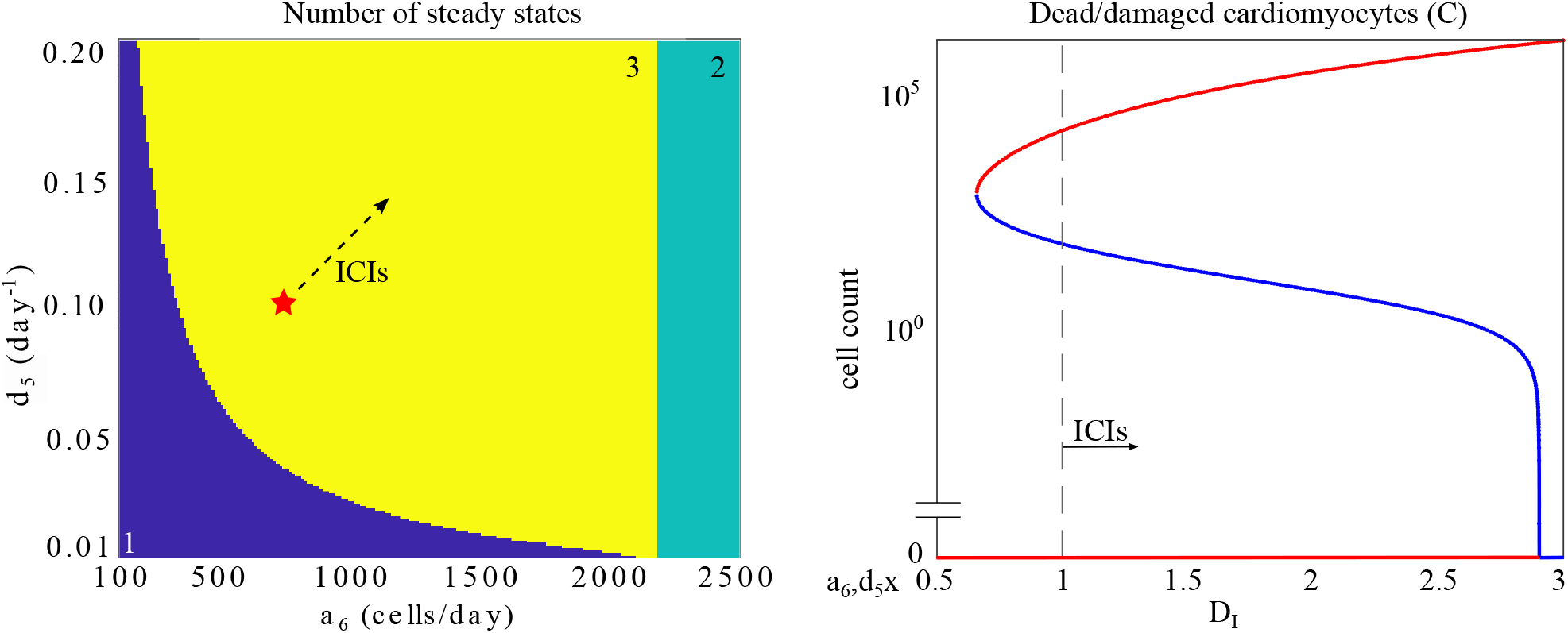
Left: Bifurcation diagram for varying values of *a*_6_(*D*_*N*_, *D*_*I*_) and *d*_5_(*D*_*I*_) as all other parameters are kept at default values. The yellow shading indicates regions of parameter space where the mode has three steady states. The green shading indicates regions with two steady states, and the purple shading indicates regions with one steady state. The red star indicates the default values of *a*_6_(*D*_*N*_, *D*_*I*_) and *d*_5_(*D*_*I*_). The dashed arrow indicates the general direction the system moves in as an increasing dose of nivolumab is applied. Right: A transcritical bifurcation occurs as the value of *a*_6_(*D*_*N*_, *D*_*I*_) is increased from its default value of 750 cells day^−1^ and the value of *d*_5_(*D*_*I*_) is increased from its default value of 0.1 day^−1^. Default values are used for all other parameters. Red indicates a stable steady state, blue indicates a saddle point. The grey dashed line indicates the default values of *a*_6_(*D*_*N*_, *D*_*I*_) and *d*_5_(*D*_*I*_). The dashed arrow indicates the direction the system moves in when nivolumab is applied.

To explore the type of bifurcation that occurs as the dose of ipilimumab is increased, *a*_6_(*D*_*N*_, *D*_*I*_) and *d*_5_(*D*_*I*_) are increased simultaneously while the number of steady states and their linear stability are tracked (see right panel in Fig. 7). Again, it is assumed here that the effect of ipilimumab on these parameters is identical in terms of the percentage change in parameter value as a function of ipilimumab dose. The grey dashed line indicates where *a*_6_(*D*_*N*_, *D*_*I*_) and *d*_5_(*D*_*I*_) attain their default values. As the dose of ipilimumab is increased, the system moves to the right away from the dashed line. Similar to nivolumab, increasing doses of ipilimumab initially lower the threshold for autoimmune myocarditis to develop before the system crosses a transcritical bifurcation and the origin becomes unstable, making the development of the disease almost inevitable. To reach this bifurcation, enough ipilimumab must be added to increase *a*_6_(*D*_*N*_, *D*_*I*_) to almost three times its default value.

In summary, applying ICIs in increasing doses can thus drastically change the linear stability of the steady states of the model. The presence of increasing levels of nivolumab or ipilimumab leads to a lower threshold for the development of autoimmune myocarditis as the saddle point moves towards the healthy steady state. For high enough levels of ICIs, a bifurcation occurs, leading to an unstable origin and a system in which the smallest perturbation away from the healthy steady state will lead to disease.

## 5. Patient-specific responses

Autoimmune myocarditis is a rare side-effect of treatment with ICIs, so the transition of the system towards resolution or chronic inflammation will differ between patients and per ICI dose. Ultimately, the aim of this research is to be able to predict how the levels of ICIs a patient can receive without developing autoimmune myocarditis, or how different patients might respond to the same treatment protocol. With the parameterised model presented in this paper, we can take preliminary steps towards this goal.

Fig. 8 shows model predictions for how an individual might respond to an immune perturbation given certain doses of nivolumab and ipilimumab have been administered. The initial conditions [*C*_0_, *P*_0_, *I*_0_, *R*_0_] = [0, 200, 0, 0] cells represent an immune perturbation that in an ICI-free system brings the systems close to the separatrix, but does not cause autoimmune myocarditis to develop. However, a relatively small change in the position of the separatrix due to the dosing of ICIs could change the perturbation outcome, so the system is sensitive to changes in drug dose for these initial conditions. For a patient who experiences fluctuations in immune cell counts of this magnitude, it would thus be important to carefully consider the amount of ICIs administered, lest they develop autoimmune myocarditis. The left panel in Fig. 8 shows how the model evolves after an influx of pathogenic CD4+ T cells as the value of *a*_6_(*D*_*N*_, *D*_*I*_) is increased step-wise up to 150% of its default value, and *d*_4_(*D*_*N*_) is simultaneously decreased up to 67% of its default value. For small doses of nivolumab (bottom three, blue time traces), the model predicts that the patient can control the immune insult and autoimmune myocarditis does not develop. For larger doses of nivolumab, however, the model predicts that the perturbation pushes the dynamics across the separatrix and disease develops (top three, green, yellow and orange time traces). The right panel in Fig. 8 shows similar results for ipilimumab, where doses of ipilimumab are represented by step-wise increases of *a*_6_(*D*_*N*_, *D*_*I*_) and *d*_5_(*D*_*I*_) up to 150% of their default values. These results demonstrate that a model such as the one presented here can aid in determining patient-specific maximum-tolerated ICI doses, taking into account the risk a specific patient runs of developing autoimmune myocarditis.

**Figure 8:**
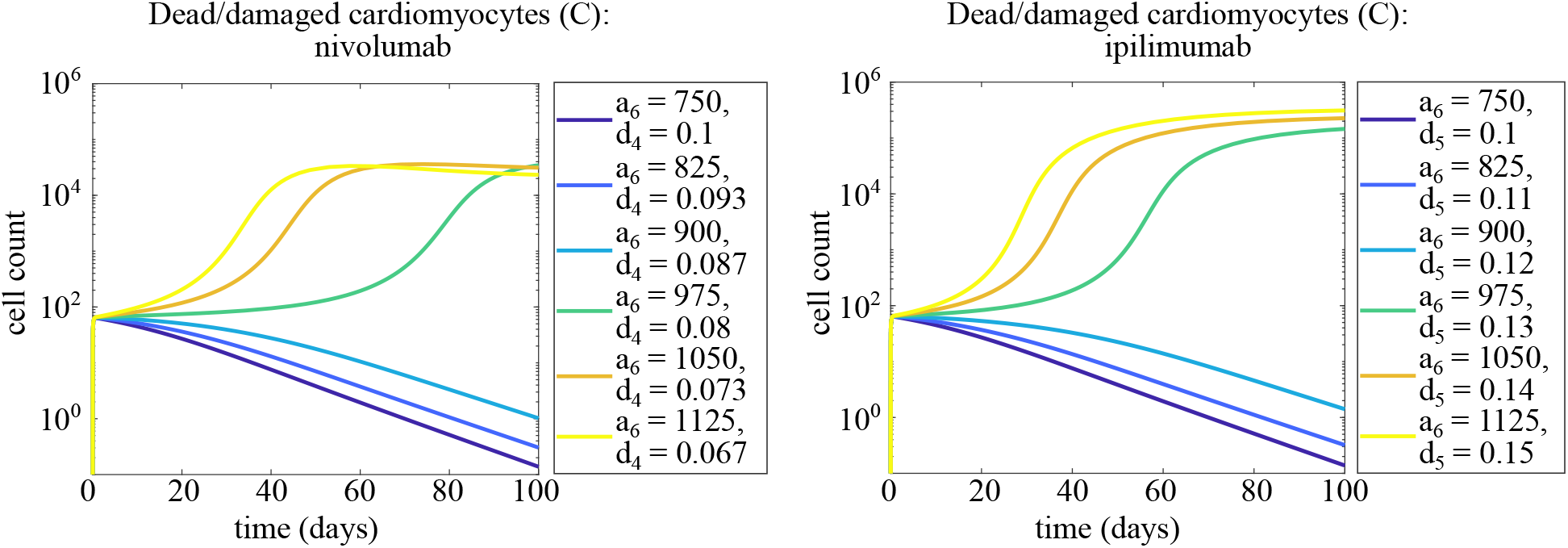
Left: Dynamics of dead/damaged cardiomyocytes in response to increasing levels of nivolumab, represented by changes in the values of *a*_6_(*D*_*N*_, *D*_*I*_) and *d*_4_(*D*_*N*_). The default parameter set is used, unless otherwise stated. Initial conditions [*C*_0_, *P*_0_, *I*_0_, *R*_0_] = [0, 200, 0, 0] cells. Right: Dynamics of dead/damaged cardiomyocytes in response to increasing levels ipilimumab, represented by changes in the values of *a*_6_(*D*_*N*_, *D*_*I*_) and *d*_5_(*D*_*I*_). The default parameter set is used, unless otherwise stated. Initial conditions [*C*_0_, *P*_0_, *I*_0_, *R*_0_] = [0, 200, 0, 0] cells.

Another important question to ask is how different patients might respond to the same dose of ICIs. To explore this, the value of *a*_1_, the rate with which pathogenic CD4+ T cells damage or kill cardiomyocytes, is varied around its default value of 80 day^−1^. The rate at which different immune processes occur is likely to vary between individuals, thus exploring variations in parameter values and the effect this has on outcome in ICI-free and ICI-dosed systems is important. Parameter *a*_1_ is chosen to illustrate the impact of individual variation. The four panels in Fig. 9 show how a slight increase or decrease in the value of *a*_1_ can have a large effect on the outcome after immune perturbation. The initial conditions [*C*_0_, *P*_0_, *I*_0_, *R*_0_] = [0, 200, 0, 0] cells are used to simulate a small immune perturbation which, in an ICI-free environment, pushes the system close to the separatrix but not across it. The top-left panel in Fig. 9 shows that in an ICI-free environment the outcome is always resolution, regardless of small changes to the value of *a*_1_. ICI doses are simulated by a 10% increase or decrease in the appropriate parameter values. This ensures that although the threshold for developing autoimmune myocarditis is lower than in an ICI-free environment, the system has not crossed the transcritical bifurcation that occurs for high doses of ICIs, so the perturbation away from the healthy steady state introduced by the initial conditions is not guaranteed to lead to the development of disease. For the simulation of combination therapy *a*_6_(*D*_*N*_, *D*_*I*_) is increased by 21% to take into account the effects of both ICIs, which are assumed to independently affect *a*_6_. The bottom left panel in Fig. 9 shows that the model predicts that administering ipilimumab does not change the outcome of inflammation and the system still resolves inflammation for all values of *a*_1_. When administering nivolumab (top right panel), however, this changes as for *a*_1_ = 85 day^−1^ the model predicts that autoimmune myocarditis will develop. When administering combination therapy (bottom right panel), all values of *a*_1_ explored here lead to autoimmune myocarditis. A small change in a parameter can thus lead to different outcomes when ICIs are applied, showing that individual patient characteristics are important to consider when it comes to designing treatment regimen.

**Figure 9:**
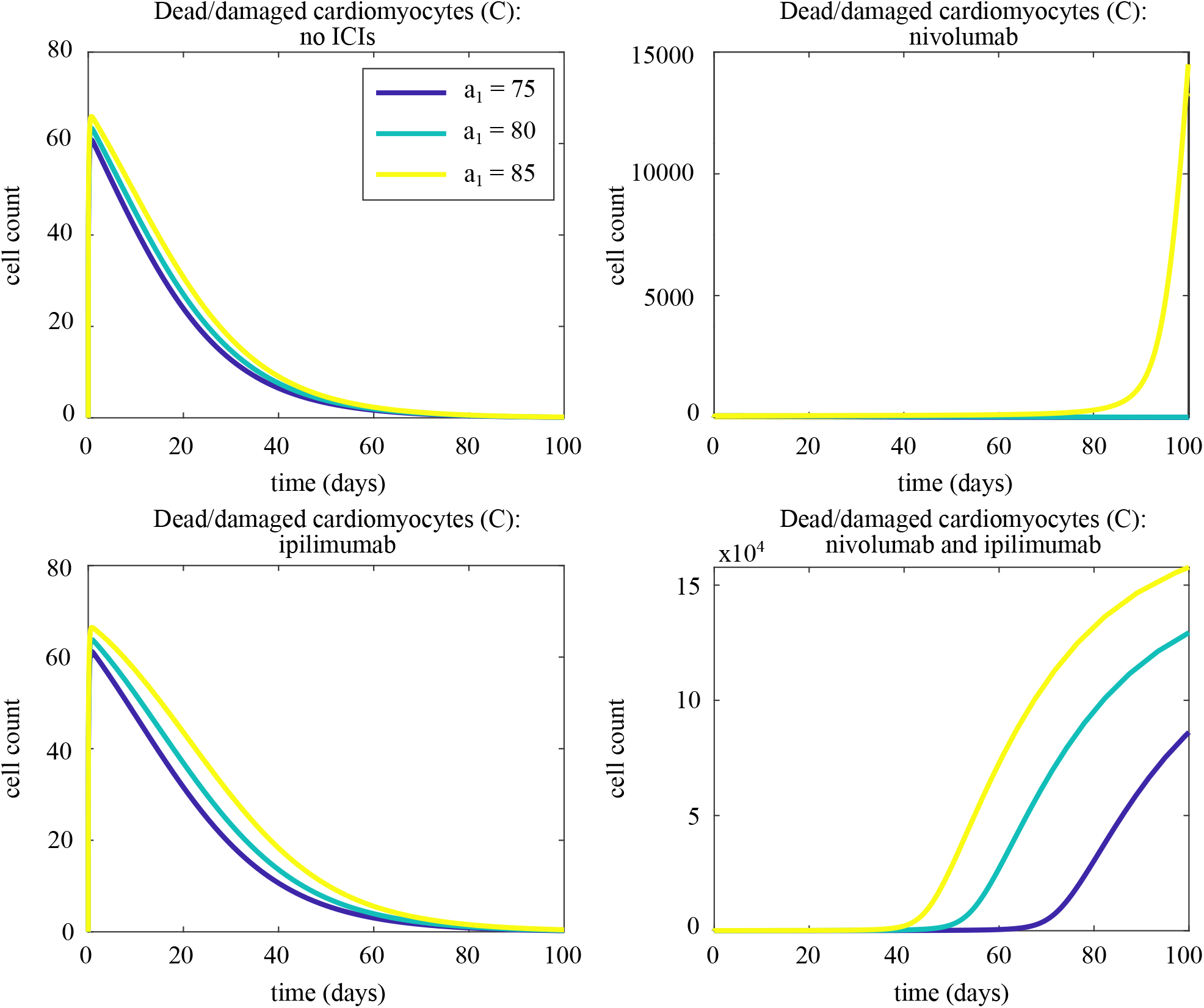
Varying outcomes for different values of *a*_1_ to a dose of nivolumab (top right), ipilimumab (bottom left), or both (bottom right), as indicated by the number of damaged/dead cardiomyocytes. ICI doses are modelled by a 10% increase in the value of *a*_6_(*D*_*N*_, *D*_*I*_) and a 10% decrease in the value of *d*_4_(*D*_*N*_) (nivolumab monotherapy), a 10% increase in the values of *a*_6_(*D*_*N*_, *D*_*I*_) and *d*_5_(*D*_*I*_) (ipilimumab monotherapy), or a 21% increase in *a*_6_(*D*_*N*_, *D*_*I*_), a 10% decrease in *d*_4_(*D*_*N*_) and a 10% increase in *d*_5_(*D*_*I*_) (combination therapy). Initial conditions [*C*_0_, *P*_0_, *I*_0_, *R*_0_] = [0, 200, 0, 0] cells, and default parameter set, unless otherwise noted.

## 6. Discussion

In this paper, we have presented the first model of autoimmune myocarditis and the effects of ICIs on its development and progression. The model describes how the numbers of damaged/dead cardiomyocytes, innate immune cells, pathogenic CD4+ T cells and regulatory T cells evolve over time, eventually leading to either resolution or chronic inflammation. The cell types and interactions in this model were included because they have been previously identified through experimental research as playing a role in the immunological mechanisms underlying this disease. The fact that the model can reproduce observations in patients with autoimmune myocarditis supports that the cell types included in this model form a required set for the model to be biologically realistic.

We have shown that the addition of ICIs makes it more likely that autoimmune myocarditis develops, i.e. a smaller perturbation away from the healthy state is required to trigger the disease. Furthermore, the number of immune cells at the disease state is higher when ICIs are present, suggesting more severe disease. We have also shown that small changes in parameter values, representing slight differences between individual patients, can influence whether a particular individual experiences resolution or chronic inflammation. For a certain ICI dose or immunological perturbation, the model predicts that some patients may be pushed towards chronic inflammation while others return to a healthy state.

Our model predicts that for a large enough dose of either nivolumab or ipilimumab, the development of autoimmune myocarditis is inevitable for even the smallest perturbation away from the healthy steady state. This is a prediction that could potentially be tested in an *in vitro* assay where the dosage of ICIs and the size of the perturbation, i.e. the number of active pathogenic CD4+ T cells added to the system, can be carefully controlled. Results of such an experiment could be used to validate the mechanisms proposed in this paper.

We highlight that the parameter values relevant for this model are highly uncertain. No data with which to calibrate the model is currently available, and the grouping of many cell types and interactions into single variables and terms makes it difficult to estimate parameter values from existing literature. As the magnitude of the perturbation away from the healthy steady state significantly impacts the timescale on which the cell counts evolve, and the timescale on which pathogenic T cells are activated is one of the criteria around which the model is parameterized, the parameter values resulting from the method we have employed are clearly dependent on the size of the perturbation. Any change to the assumed magnitude of the perturbation may require changes to the parameter values to ensure that all criteria are still met. However, we highlight that the qualitative behaviours of the system are not affected.

Should patient-specific data with which to calibrate become available, this model could be used to estimate the maximum tolerated dose of ICIs. Fitting the model parameters to patient-specific data allows us to compute where the saddle point lies, how far away it is from the healthy steady state and thus how big a perturbation away from the healthy steady state a patient can tolerate. This enables assessment of the risk of a patient developing autoimmune myocarditis for a certain dose of ICIs. It also facilitates predictions about outcomes for individual patients and the timeline on which disease may develop. This is all useful data that could potentially inform the treatment plan for specific patients.

To be able to have confidence in model predictions, we must ensure that the parameters of the model are identifiable given the available data. To obtain an understanding of the type and quality of data required to establish parameter values for this model, we conducted a parameter identifiability analysis. Structural identifiability analysis performed using DAISY indicated that the model is only structurally identifiable if all four variables are measured as outputs. To assess the practical identifiability of the model parameters, the maximum loglikelihood profiles of subsets of the 13 parameters in the ICI-free model are computed, using the methods explained by Simpson et al. [77]. An example of the results of the practical identifiability analysis are shown in Fig. 10, where the profile loglikelihoods show that the death rates and *ϕ* form a practically identifiable subset of parameter space. More details regarding the methods and results of the practical identifiability analysis can be found in Appendix A.

**Figure 10:**
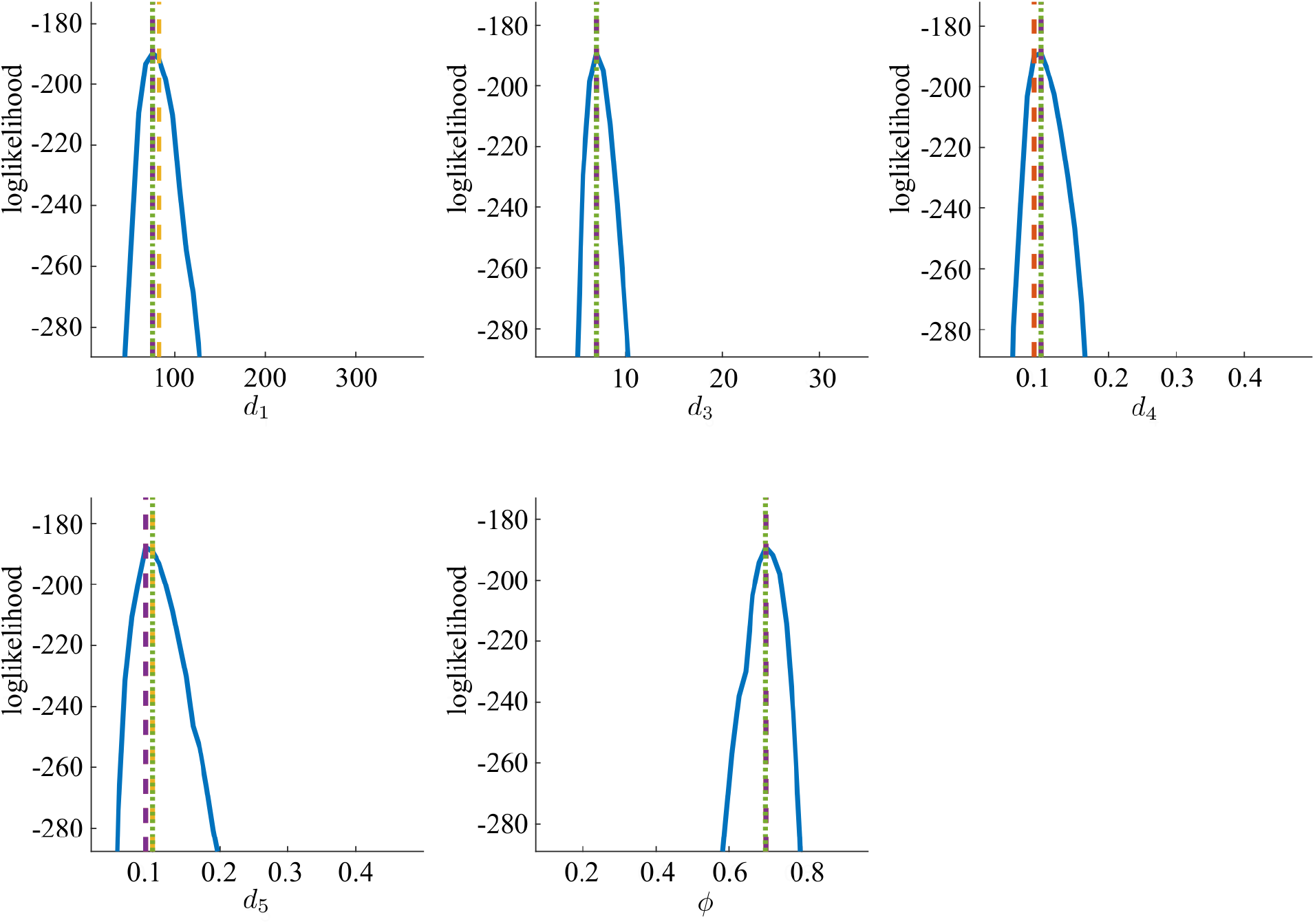
Maximum loglikelihood profiles of the subset of parameters *d*_1_, *d*_3_, *d*_4_, *d*_5_ and *ϕ* in the ICI-free model. Details regarding the implementation of the algorithm explain by Simpson et al. [77] can be found in Appendix A.

In the future, this model may be used together with methods from optimal control theory to predict optimal drug regimen. If the goal is to maximally attack the cancer with ICIs while preventing the development of autoimmune myocarditis, the frequency and quantity of ICI dosage needs to be carefully planned. Our results demonstrate that as the drug dose increases, the saddle point approaches the healthy steady state, which leaves less and less room for fluctuations in the number of DCs and pathogenic immune cells without crossing over into the basin of attraction of the disease steady state. Optimal control theory can be used to determine where the ideal balance between the opposing aims of treating the tumour and preventing autoimmunity lies.

With this model, we have taken the first steps in consolidating the current knowledge of autoimmune myocarditis and the effects of ICIs on its development and progression. It will be important to further investigate which cell types and molecules are the critical players in this disease in order to work towards prevention of this side effect of ICIs. Though much work remains to be done, we have started on the path towards a mathematical model of autoimmune myocarditis and the role of ICIs that can be used to determine which cell types, cytokines and other components are critical in the development and progression of this severe side effect. With these essential immune components identified, work can start towards reducing the risk of severe cardiotoxic side effects of ICIs in patients.

## Conflict of Interest Statement

The authors declare that the research was conducted in the absence of any commercial or financial relationships that could be construed as a potential conflict of interest.

## Author Contributions

S.A. van der Vegt prepared and edited the manuscript. Y.-J. Wang, L. Polonchuk, K. Wang, S.L. Waters and R.E. Baker edited the manuscript. All authors designed the research.

## Acknowledgments

The authors would like to thank Prof. Mark Coles for helpful discussion and suggestions. S.A.V. is supported by the EPSRC (EP/L016044/1). R.E.B is a Royal Society Wolfson Research Merit Award holder.

## Appendix A. Practical identifiability analysis

The large number of parameters and relatively small number of variables in the model makes a comprehensive practical identifiability analysis infeasible. The parameter set is therefore split into subsets to investigate which subsets of model parameters are identifiable. We use the profile loglikelihood method to explore the practical identifiability of parameters [77]. In this method, the parameter of interested is varied over a set range and for each value, the values of the other parameters in the subset are optimized to maximize the loglikelihood.

A synthetic data set is obtained by simulating the model for 60 days, obtaining an “observation” of all four variables on days 0, 1, 2, 5, 10, 30 and 60. Structural identifiability analysis performed with DAISY indicates that the model parameters are only structurally identifiable when all variables are observed, therefore observing only a subset of the variables will guarantee practical unidentifiability. The observations are concentrated in the first few days so the fast early dynamics of the system are captured. Normally distributed measurement noise with a standard deviation of 5% of the cell count observed is added to all data points.

The initial conditions are [*C*_0_, *I*_0_, *P*_0_, *R*_0_] = [0, 0, 10^4^, 0] to ensure the model is identifiable for scenarios where disease develops. A representative synthetic data set is shown in Fig. A.11.

**Figure A.11:**
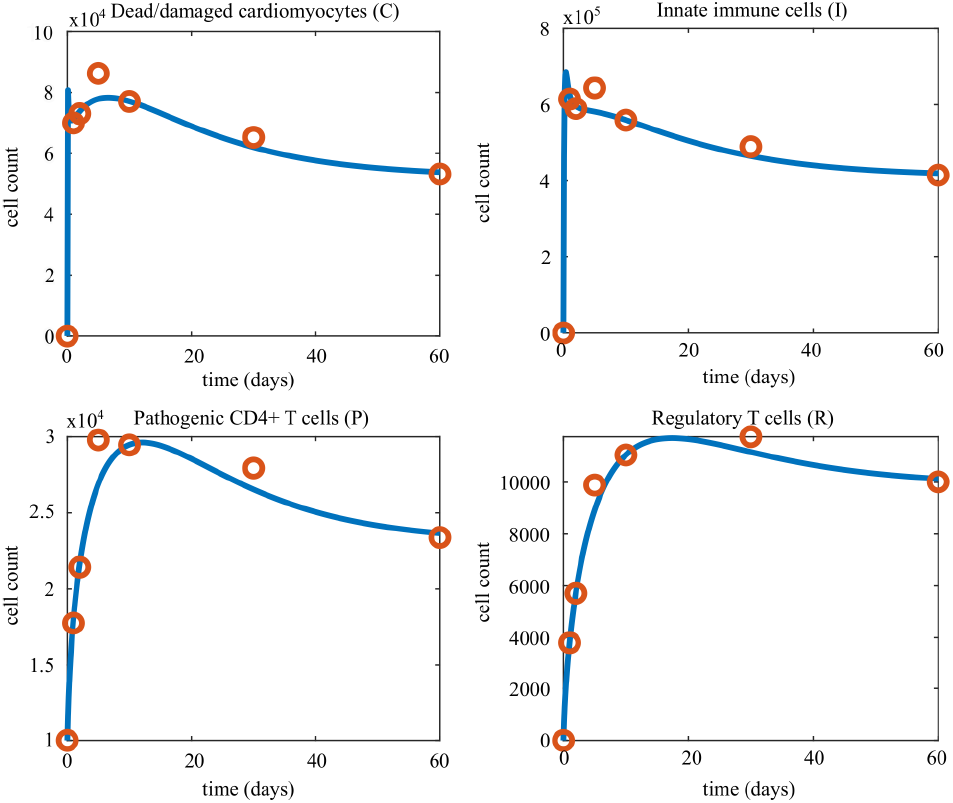
A representative simulation (blue line) and resulting synthetic data set (red circles). Observations for the data set are made on 0, 1, 2, 5, 10, 30 and 60, after which normally distributed noise with a standard deviation of 5% of the cell count is added to each data point.

Parameters are varied between 0.1 and five times their default values, in steps of 0.1. During the maximization of the loglikelihood, the lower bound for all parameters to be optimized is 10^−14^. The upper bound is infinity for all parameters except for *ϕ*, which cannot exceed one.

We used globalsearch to maximize the loglikelihood, with the optimizer fmincon. For the first value of the parameter of interest (i.e. 0.1 times its default value), the initial guesses for the values of the parameters to be optimized are their respective default values. For the subsequent values of the parameter of interest, the optimized values of the other parameters from the previous step are used as initial guesses. For fmincon, we use the algorithm spq. We further use FunctionTolerance and XTolerance 10^−10^. The function to be minimized is the loglikelihood squared, computed as

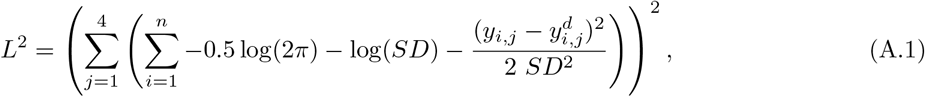

where *y*_*j*_ is the model output of the variable *j* for a given parameter set, *y*^*d*^ is the data set, *i* denotes one of *n* data points, and *SD* is the standard deviation for a specific data point, computed as 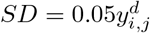.

The results in Fig. 10 show that the death rates and *ϕ* form a practically identifiable subset with the synthetic data set created with the above requirements. The set of the activation rates, however, do not form a practically identifiable subset given the specific settings used here, as is shown in Fig. A.12 where a subset of the activation rates consisting of the parameters *a*_1_, *a*_2_, *a*_4_, and *a*_5_ is shown to be practically unidentifiable.

**Figure A.12:**
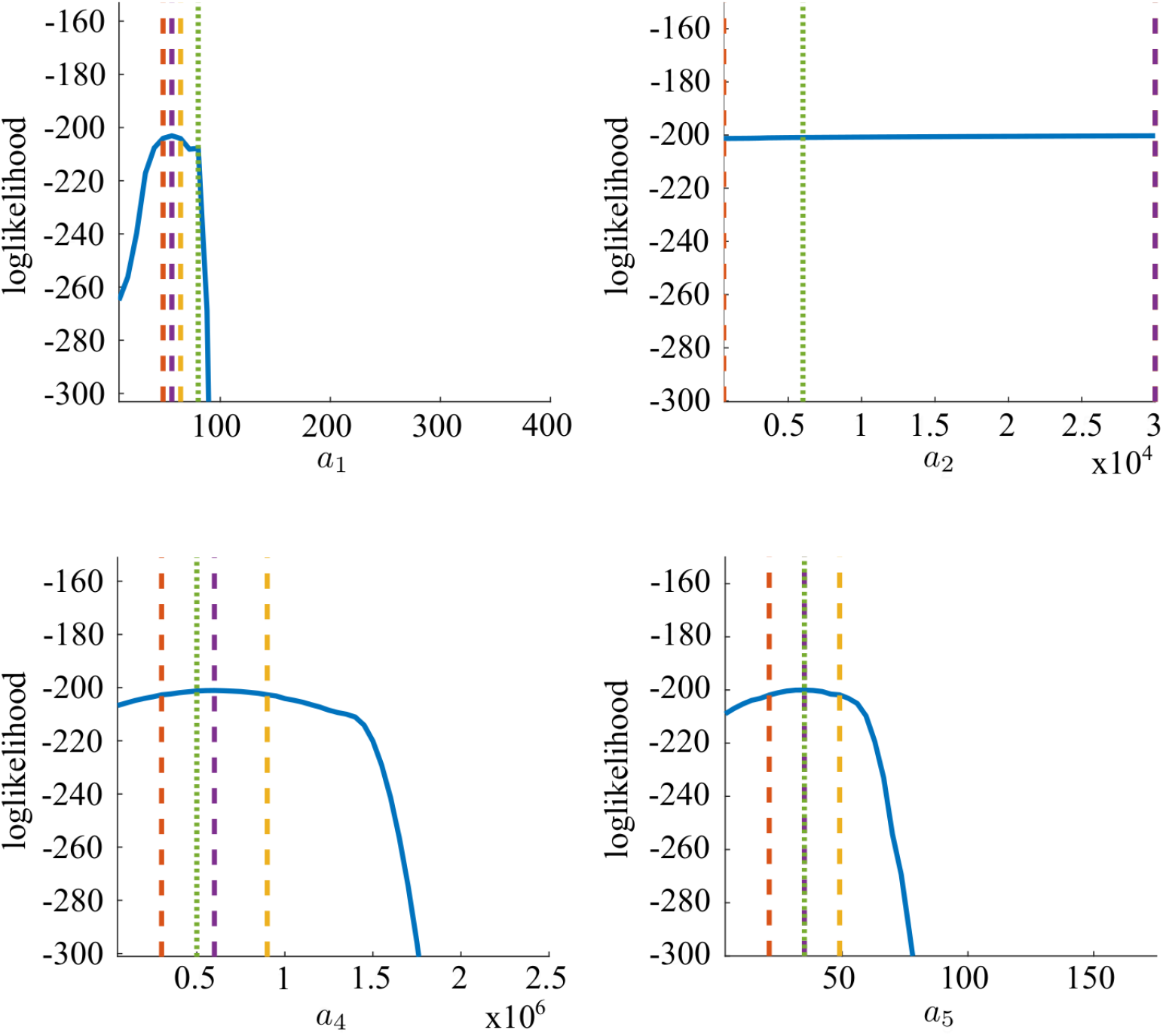
Profile loglikelihoods for the parameter subset *a*_1_, *a*_2_, *a*_4_ and *a*_5_, computed according to the method described in [77]. Green dotted line indicates the true value of the parameter. Purple dashed line indicates the value of the paramter for which the maximum loglikelihood value is attained. The red and yellow dashed lines indicate the 95% confidence interval around the loglikelihood maximum.

The subsets can also be formed by grouping parameters by ODE they appear in. As is to be expected, this leads to practically unidentifiable subsets as parameter values in these subsets can be adjusted to compensate for the varying value of the parameter of interest. An example of this is shown in Fig. A.13, which shows the flat profile loglikelihoods for the parameter subset *a*_1_, *a*_2_ and *d*_1_, all of which appear in the ODE describing the dynamics of damaged/dead cardiomyocytes.

**Figure A.13:**
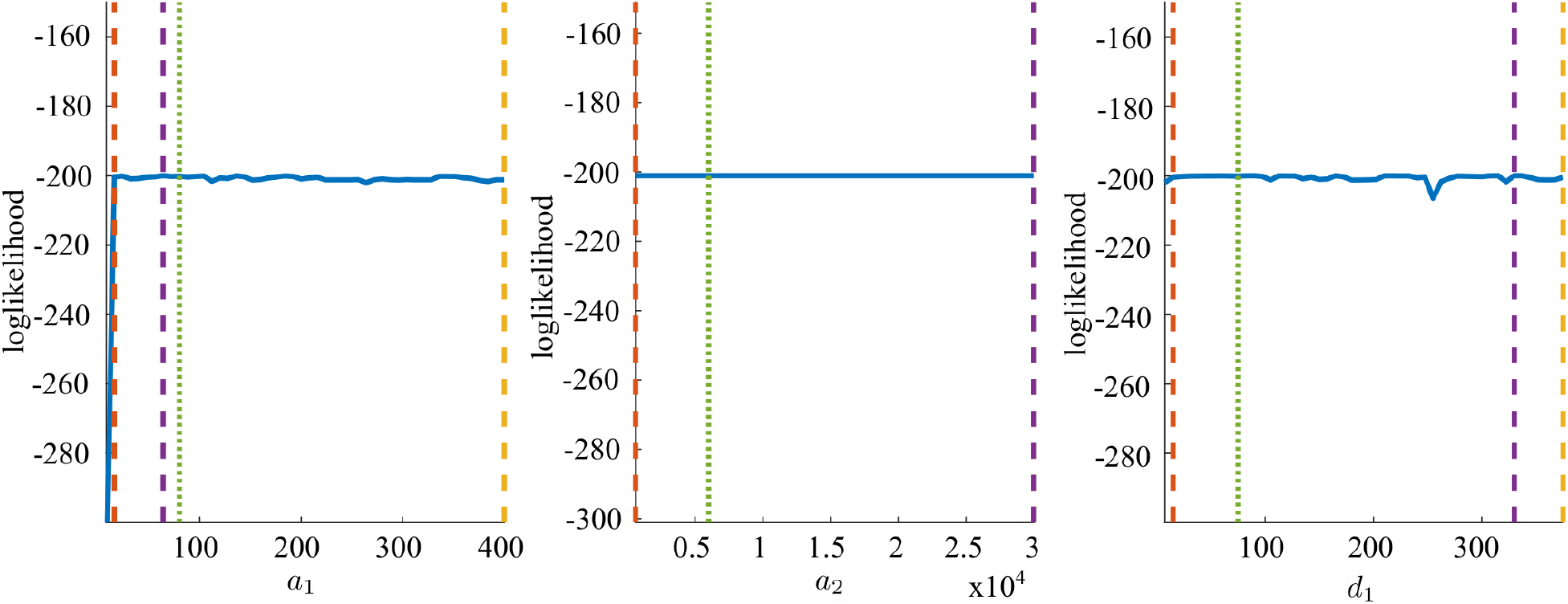
Profile loglikelihoods for the parameter subset *a*_1_, *a*_2_ and *d*_1_, computed according to the method described in [77]. Green dotted line indicates the true value of the parameter. Purple dashed line indicates the value of the parameter for which the maximum loglikelihood value is attained. The red and yellow dashed lines indicate the 95% confidence interval around the loglikelihood maximum.

## Notes

### Competing Interest Statement

The authors have declared no competing interest.

